# PIF transcription factors link a neighbor threat cue to accelerated reproduction in Arabidopsis

**DOI:** 10.1101/377663

**Authors:** V.C. Galvāo, A.S Fiorucci, M. Trevisan, J.M. Franco-Zorrilla, A. Goyal, E. Schmid-Siegert, R. Solano, C. Fankhauser

## Abstract

Changes in light quality indicative of competition for this essential resource influence plant growth and developmental transitions. Little is known about neighbor proximity-induced acceleration of reproduction. phytochrome B (phyB) senses light cues from plant competitors ultimately leading to the expression of the floral inducers *FLOWERING LOCUS* (*FT*) and *TWIN SISTER of FT* (*TSF*). Here we show that three PHYTOCHROME INTERACTING FACTOR (PIF) transcriptional regulators act directly downstream of phyB to promote expression of *FT* and *TSF*. Neighbor proximity enhances PIF accumulation towards the end of the day coinciding with enhanced floral inducer expression. We present evidence for direct PIF-mediated *TSF* expression. The relevance of our findings is illustrated by the prior identification of *FT*, *TSF* and *PIF4* as loci underlying flowering time regulation in nature.

**One Sentence Summary:** PIF transcription factors mediate reproductive transition in response to neighbor proximity light cues in Arabidopsis.

---

Plants depend on sunlight to fuel photosynthesis. Therefore, growing with potentially reduced light availability, as encountered in dense plant communities, constitutes a threat for plant growth and development. Plants perceive potential competitors because of the reflected far-red (FR) light from neighbors, resulting in reduced red (R)/FR ratio (R/FR), which leads to the conversion phytochromes (phy) photoreceptors to their inactive Pr form. In shade-intolerant plants, this triggers organ elongation to outgrowth competitors and precocious flowering (*1*). Accelerated flowering results from the transcriptional induction of the florigen *FLOWERING LOCUS T* (*FT*) and its close homologue *TWIN SYSTER OF FT* (*TSF*) in the vasculature (*2-5*), followed by their transport to the shoot apical meristem. In Arabidopsis, low R/FR ratio promotes floral transition in a photoperiod-dependent manner (*6*), in agreement with the attenuated low R/FR response of the photoperiodic mutant *constans* (*co*) (*2, 6*). PHYTOCHROME AND FLOWERING TIME 1 (PFT1) was proposed to control flowering in response to simulated shade (*4*), but was later shown to respond normally to low R/FR (*6*). Here we investigate how phytochromes mediate early flowering in response to low R/FR.

The bHLH transcription factors PHYTOCHROME-INTERACTING FACTORS (PIF) play major roles in neighbor detection responses downstream of phyB (*7, 8*). Enhanced *PIF* expression induces precocious flowering through *FT* and *TSF* in the phloem (*9-11*). Moreover, plants with impaired HFR1 function, a repressor of PIF activity (*12*), display an increased *FT* expression in response to low R/FR (*13*). Therefore, we hypothesized that PIFs might control flowering time in response to low R/FR. We scored the flowering transition of PIF loss-of-function mutants under simulated neighbor detection (fig. S1; hereafter referred to as low R/FR) and showed that PIF7 plays a prominent function in flowering transition under low R/FR (Table 1, experiment 1). In addition, mutations in *PIF4* and *PIF5* further enhanced the *pif7* phenotype, indicating that these genes also contribute to the response (Fig. 1A; Table 1, experiment 1, significant interaction between genotype and condition p < 0.001; Fig. S2). Moreover, while *pif3pif4pif5* and *pif4pif5pif7* both flower slightly late in high R/FR, specifically *pif4pif5pif7* flowered later than the wild type in low R/FR (Fig. S2). Next, we checked whether PIFs mediate precocious flowering of the constitutive shade-avoidance mutant *phyB*. Consistent with our data in low R/FR, mutations in *PIF4, PIF5* and *PIF7* were required to fully suppress early flowering in *phyB*, including in non-inductive photoperiods (Fig. 1B; Table 1, experiment 2; Fig. S3). We conclude that *PIF4*, *PIF5* and *PIF7* act genetically downstream of phyB to control low R/FR-induced flowering.

**Fig. 1.**
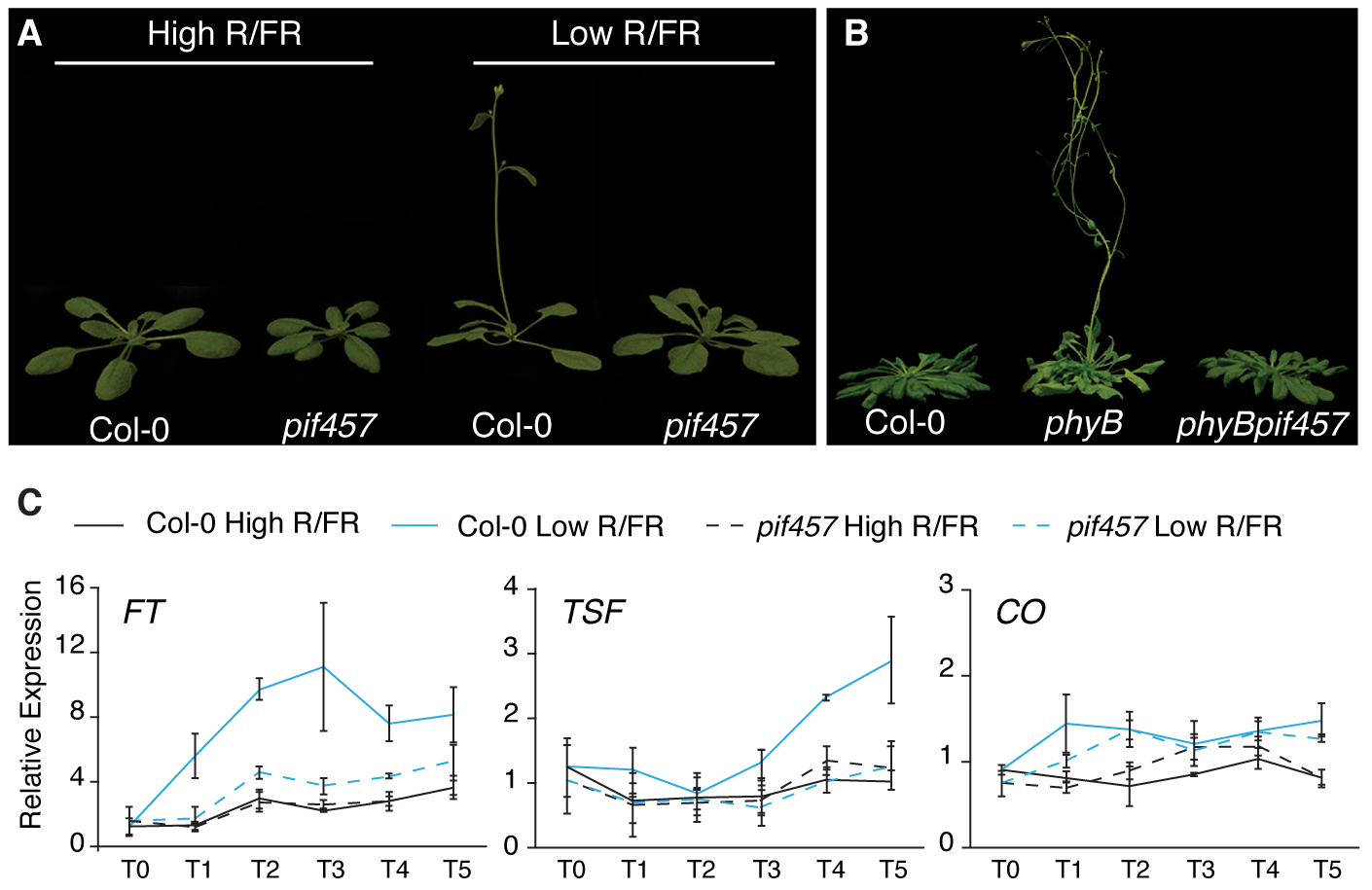
PIFs mediate flowering in low R/FR downstream of phyB. (A) Phenotype of 22-day-old Col-0 and *pif4pif5pif7* grown under LD at 22°C in high and low R/FR and (B) 53-day-old Col-0, *phyB* and *phyBpif4pif5pif7* under SD at 22°C in high R/FR. (C) *FT, TSF and CO* mRNA expression following a shift from high to low R/FR in Col-0 and *pif4pif5pif7*. Plants were grown for 5 days under LD at 22°C in high R/FR and samples were harvested at ZT 15-16 before (T0) and 1, 2, 3, 5, and 7 days after transfer to low R/FR.

**Table 1.** Flowering time of mutant lines represented as leaf number and days to flowering. RLN, rosette leaf number; CLN, cauline leaf number; TLN, total leaf number; SD, standard deviation; n, number of individuals; high, high R/FR; low, low R/FR. Superscript letters represent the significance groups at p-value < 0.01 using ANOVA followed by *post hoc* Tukey HSD test.

| Genotype | Light | RLN | CLN | TLN | SD | Days | SD | n |
| --- | --- | --- | --- | --- | --- | --- | --- | --- |
| **Experiment 1 - LD** |  |  |  |  |  |  |  |  |
| Col-0 | High | 10.5 | 2.9 | 13.4<sup>c</sup> | 0.9 | 22.3<sup>b</sup> | 1.0 | 30 |
|  | Low | 4.9 | 2.2 | 7.1<sup>f</sup> | 0.7 | 12.7<sup>e</sup> | 1.3 | 31 |
| *pif7* | High | 12.5 | 2.9 | 15.5<sup>a</sup> | 1.3 | 24.1<sup>a</sup> | 1.0 | 18 |
|  | Low | 7.2 | 2.9 | 10.1<sup>e</sup> | 1.0 | 18.1<sup>c</sup> | 1.4 | 23 |
| *pif4 pif5* | High | 11.2 | 2.9 | 14.1<sup>bc</sup> | 1.0 | 23.8<sup>a</sup> | 0.9 | 27 |
|  | Low | 5.4 | 2.3 | 7.6<sup>f</sup> | 0.6 | 14.7<sup>d</sup> | 1.7 | 31 |
| *pif4 pif5 pif7* | High | 12.3 | 2.5 | 14.8<sup>ab</sup> | 1.1 | 24.2<sup>a</sup> | 0.7 | 22 |
|  | Low | 8.6 | 2.7 | 11.3<sup>e</sup> | 0.9 | 21.6<sup>b</sup> | 1.7 | 28 |
| **Experiment 2 - SD** |  |  |  |  |  |  |  |  |
| Col-0 | High | 60.2 | 8.5 | 68.7<sup>a</sup> | 5.9 | 60.2<sup>a</sup> | 3.3 | 27 |
| *phyB* | High | 19.4 | 4.6 | 24<sup>e</sup> | 3.3 | 36.2<sup>d</sup> | 2.3 | 33 |
| *phyB pif7* | High | 35.9 | 6.1 | 42<sup>c</sup> | 4.9 | 46.2<sup>c</sup> | 2.9 | 30 |
| *phyB pif4 pif5* | High | 46.9 | 8.8 | 55.7<sup>b</sup> | 4.1 | 50.2<sup>b</sup> | 2.5 | 29 |
| *phyB pif4 pif5 pif7* | High | 59.1 | 8.2 | 67.3<sup>a</sup> | 4.0 | 61.1<sup>a</sup> | 3.5 | 33 |
| **Experiment 3 - LD** |  |  |  |  |  |  |  |  |
| *ft* | High | 42.9 | 7.9 | 50.7<sup>c</sup> | 3.7 | 47.4<sup>c</sup> | 3.3 | 16 |
|  | Low | 23.9 | 6.5 | 30.4<sup>d</sup> | 2.0 | 33.4<sup>d</sup> | 2.2 | 17 |
| *ft tsf* | High | 60.2 | 10.6 | 70.8<sup>a</sup> | 3.6 | 54.9<sup>b</sup> | 3.8 | 14 |
|  | Low | 48.2 | 8.8 | 57<sup>b</sup> | 2.6 | 46.9<sup>c</sup> | 1.8 | 15 |
| *ft pif4 pif5 pif7* | High | 52.5 | 7.9 | 60.4<sup>b</sup> | 8.8 | 62.9<sup>a</sup> | 12.2 | 13 |
|  | Low | 39.9 | 7.5 | 47.4<sup>c</sup> | 2.7 | 45.6<sup>c</sup> | 1.3 | 15 |
| **Experiment 4 - LD** |  |  |  |  |  |  |  |  |
| Col-0 | High | 10.5 | 3.2 | 13.7<sup>d</sup> | 1.1 | 19.7<sup>e</sup> | 1.6 | 19 |
|  | Low | 5.5 | 2.3 | 7.9<sup>e</sup> | 0.5 | 12.5<sup>f</sup> | 0.9 | 24 |
| *co* | High | 46.2 | 5.8 | 52.1<sup>b</sup> | 2.8 | 43.4<sup>c</sup> | 2.0 | 20 |
|  | Low | 36.7 | 7.2 | 44<sup>c</sup> | 2.4 | 37.6<sup>d</sup> | 1.6 | 24 |
| *co pif4 pif5 pif7* | High | 59.1 | 7.6 | 66.7<sup>a</sup> | 4.2 | 60.9<sup>a</sup> | 5.0 | 19 |
|  | Low | 47.5 | 6.9 | 54.4<sup>b</sup> | 2.1 | 47.4<sup>b</sup> | 3.1 | 24 |

Because flowering in low R/FR depends on the growth condition and genetic background (*2, 6, 14*), we tested the flowering response of *ft*, *tsf, fttsf* and *co*. In our conditions *ft* and *tsf* mutants responded strongly to low R/FR (*2*), and *phyBtsf* flowered as *phyB* indicating that neither FT nor TSF alone accounted for early flowering (Fig. S4 and S5). In contrast, *fttsf* double mutant presented a reduced low R/FR response, similar to *co* (Fig. S4), confirming that FT and TSF together are required to accelerate flowering in low R/FR (*2*). Next, we determined whether PIFs contribute to *FT* and *TSF* transcriptional regulation in low R/FR (*2, 4-6*). Transcriptome data (*15*) showed that *FT* mRNA levels increased in cotyledons within 90 minutes after transfer to low R/FR, while such a rapid induction was not observed for *TSF* (Fig. S6). We therefore monitored *FT* and *TSF* expression in the wild type and *pif4pif5pif7* for several days after transfer from high to low R/FR at ZT16, as *FT* and *TSF* expression peak at dusk (*6, 16*). *FT* and *TSF* expression were similar in *pif4pif5pif7* and wild-type plants in high R/FR. In contrast *FT* and *TSF* up-regulation by low R/FR were strongly impaired in *pif4pif5pif7* (Fig. 1C). Consistent with the importance of PIF-dependent *TSF* up-regulation, *ftpif4pif5pif7* quadruple mutants flowered later compared to *ft* and similar to *fttsf* under low R/FR (Table 1, experiment 3; Fig. S7). In contrast, *CO* mRNA expression was only marginally increased by light treatments in both genotypes (Fig. 1C). These results identify PIFs as important mediators of FT and TSF-induced early flowering in response to low R/FR.

To better understand how PIFs control *FT* and *TSF* expression we investigated their temporal and spatial expression pattern. Consistent with the vascular expression of *FT* and *TSF* during floral transition (*2, 17*), promoter-GUS fusions showed broad *PIF4* and *PIF5* expression, including the leaf vasculature in seedlings and adult plants (Fig. 2A; Fig. S8). This is consistent with tissue-specific expression analysis of *PIF4*, *PIF5* and *PIF7* (*18*) indicating that *PIF4*, *PIF5*, *PIF7*, *FT* and *TSF* are all expressed in the vasculature. *FT* mRNA expression in the wild type displayed two strong peaks in response to low R/FR, the first early in the light period and the highest peak around dusk (Fig. S9A) (*6*). In contrast, there was no induction of *FT* expression by low R/FR in *pif4pif5pif7* (Fig. S9A). *TSF* expression and its regulation by low R/FR and the PIFs were very similar to *FT* (Figure S9B). *PIF7* expression showed a strong diel oscillation with a peak in the morning as previously observed for *PIF4* and *PIF5* (*19*) (Fig. S9C). However, low R/FR ratio did not affect significantly *PIF7* mRNA expression throughout the day (fig. S9C) (*15*), suggesting that transcription regulation of *PIFs* alone cannot account for increased *FT* and *TSF* expression in low R/FR. Given that phyB inactivation under low R/FR stabilizes PIF4 and PIF5 proteins (*7*) we decided to investigate PIF protein accumulation. Using genomic HA-tagged lines driven by their own promoters (Fig. S10A-D), we observed diel protein oscillation of HA-tagged PIF4, PIF5 and PIF7 matching mRNA levels (Fig 2B, C; Fig. S11). Interestingly, PIF4 and PIF5, proteins accumulated to higher levels in low R/FR specifically toward the end of the day, correlating with *FT* and *TSF* expression (Fig 2B; Fig. S9 and S11). Such a regulation was less apparent for PIF7 (Fig 2C), however PIF7 nuclear import is induced by low R/FR (*20*), indicating a different mode of low R/FR regulation for this PIF.

**Fig. 2.**
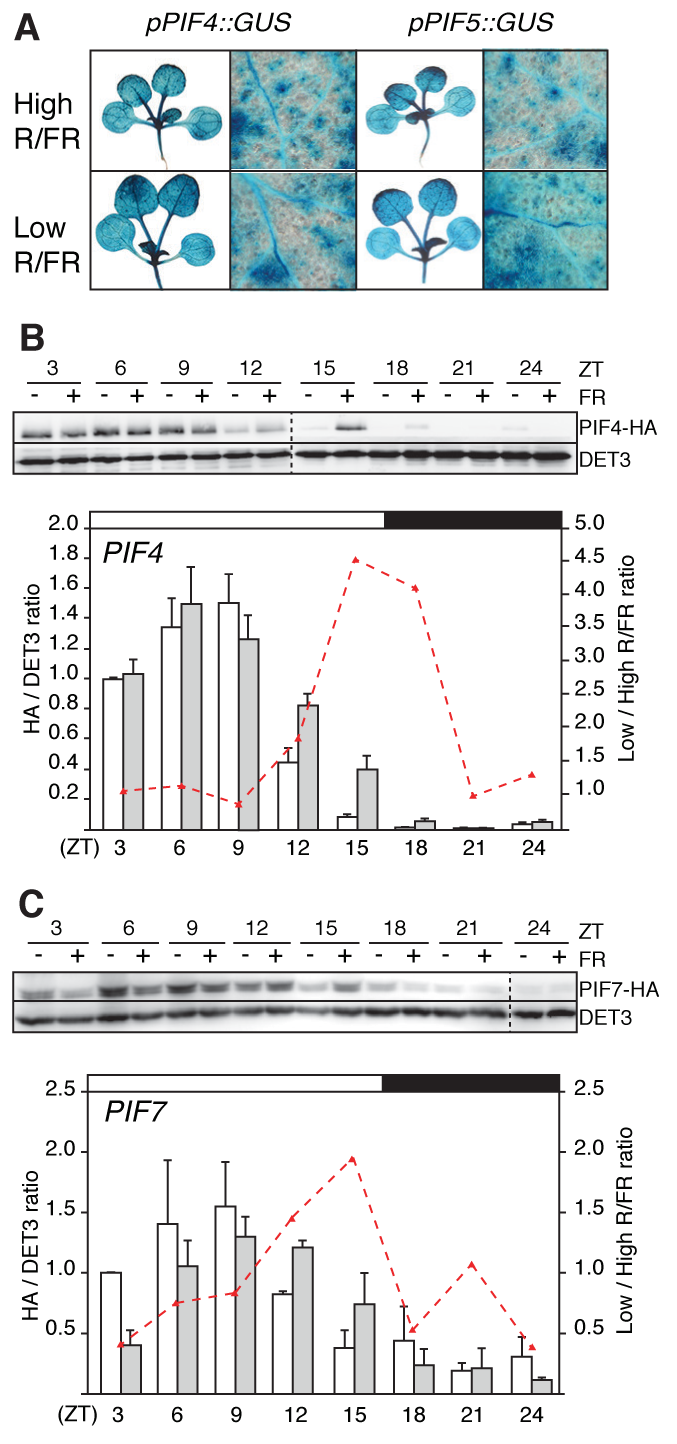
PIFs expression pattern and protein stability in high and low R/FR. (A) Representative GUS staining of *pPIF4::GUS* and *pPIF5::GUS* lines harvested 9-days after sowing at ZT16 in high and low R/FR. Protein level of (B) *pif4-101/pPIF4::PIF4-3HA* and (C) *pif7-2/pPIF7::PIF7-3HA* in 10-11-day-old plants harvested every 3 hours. White (high R/FR) and grey (low R/FR) bars correspond to the average protein levels of 3 biological replicates and at least 2 technical replicates relative to DET3. Red dashed line represent the protein level ratio of low/high R/FR. Error bars represent standard deviation and white and black bars on top of each chart represent the light and dark phases, respectively.

Because TSF was shown to integrate environmental signals to influence flowering time in nature (*21*), we focused our analysis on PIF regulation of *TSF* expression. PIF4 and PIF5 preferentially bind to G-boxes (CACGTG) and PBE-boxes (CATGTG) (*22, 23*). Because PIF7 plays a central role in low R/FR-induced flowering and little is known about its DNA binding preference, we tested its DNA binding specificity using protein-binding microarrays. In agreement with recent DAP-seq data (*24*) and similar to other PIFs (*22, 23*), we found that PIF7 binds with high affinity to a G-box (Fig 3A). Moreover, as observed for other PIFs, among E-boxes it showed the highest affinity for the PBE-box (Fig. 3A). We identified 2 PBE-boxes in the *TSF* promoter located 990 and 437 bases upstream of the ATG (Fig. 3B). Interestingly, the analysis of previously published ChIP-seq data (*25*) revealed a high-confidence PIF4 binding peak overlapping the first PBE-box (-437) (Fig. 3B). To test whether these PBE-boxes are biologically relevant for PIF-mediated *TSF* expression we fused its promoter with luciferase and performed transient expression assays in *Nicotiana benthamiana*. Consistent with PIFs directly regulating *TSF* expression, PIF4 and PIF7 led to *TSF* expression that was almost completely abolished by a single nucleotide mutation of either 1 (-437) or both PBE-boxes (Fig. 3C; fig S12). Taken together, our data suggest that PIF4 and PIF7 directly bind to PBE-boxes at *TSF* promoter to induce its expression.

**Fig. 3.**
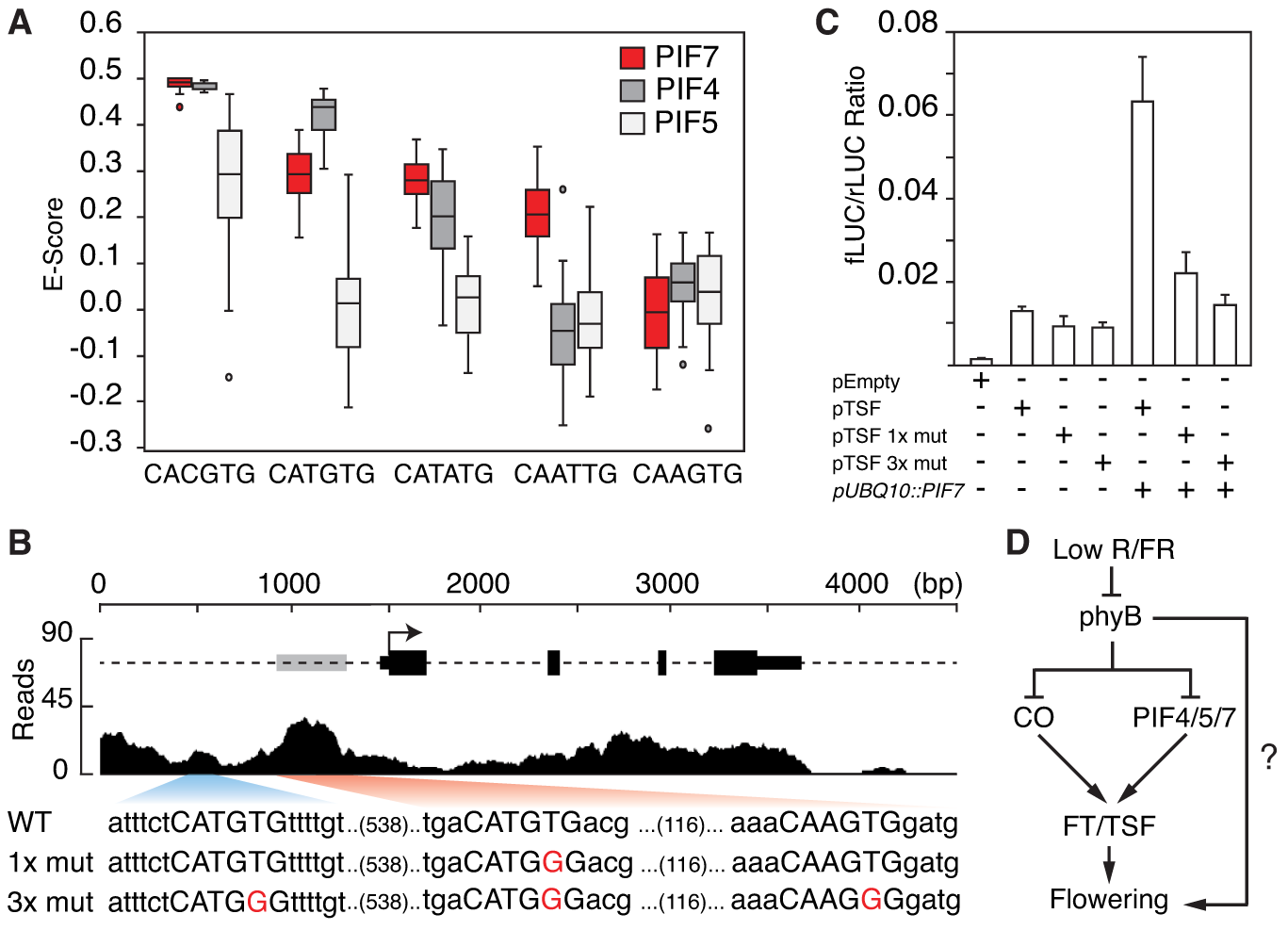
PIF proteins directly regulate *TSF* expression. (A) PIF7 preferentially bind to G-boxes and PBE-boxes in protein-binding microarray using PIF7_bHLH-MBP (Table S2). PIF4 and PIF5 data are represented for comparison (*23*). (B) Representation of PIF4 ChIP-seq reads mapped to the *TSF locus* (*25*). Grey box represent high confidence PIF4 binding peak at *TSF* promoter and nucleotide sequence represent the WT and mutant version containing 1 (1x mut) or 3 mutations (3x mut) used for transient dual-luciferase assays in *N. benthamiana* (C). Luciferase ratio corresponds to the average of *pTSF::fireflyLUC* and *p35S::renillaLUC* ratio of four independent infiltrations and error bars correspond to standard deviation. (D) Model of low R/FR-regulated flowering time.

Our experiments identify PIF4, PIF5 and PIF7 as transcription factors acting downstream of phyB to induce flowering response to neighbor proximity through the floral inducers *FT* and *TSF*. However, our genetic data indicate that additional mechanisms contribute to this regulation. Indeed, similar to *co* and *fttsf* (fig S4), *copif4pif5pif7* are still responsive to low R/FR (Table 1, experiment 4). Therefore, we identify one important flowering-time control mechanism operating in low R/FR and provide evidences for direct regulation of *TSF* by the PIFs. Interestingly, as previously shown for CO (*2*), low R/FR leads to increased PIF4 and PIF5 proteins levels (Fig. 2B-C; Fig. S11), consistent with the proposed coordinated regulation of *FT* and *TSF* expression by PIFs and CO (*11*) (Fig. 1, 2, 3C, 3D, table 1). The regulation we uncovered here is likely to be significant in natural environments as “florigen” genes *FT* and *TSF* as well as *PIF4* were identified as genes underlying regulation of flowering time in nature (*21, 26*). Understanding this regulatory mechanism may also be relevant to increase yields on restricted agricultural land.

## Acknowledgments

We thank Koji Goto for *pFT::GUS* line and *co-101* seeds, Andrea Maran and Séverine Lorrain for genomic *pPIF5::PIF5-3HA* and *pPIF4/5::GUS* lines, Markus Schmid for *ft-10* mutant, Prof. Hongtao Liu for *pGREENII-0800* vector, Rodrigo S. Reis dual-luciferase assistance, Genomic Technologies Facility (GTF) for qPCR assistance, Adriana Arongaus and Martina Legris for critical reading of the manuscript. Work in the Fankhauser lab is funded by the University of Lausanne and grants from the Swiss National Science Foundation (n° 310030B_179558 and CRSII3_154438). VCG was supported by EMBO long-term fellowship (ALTF 293-2013).

